# Library Size in Spatial ATAC-seq: Technical Confounder or Biology?

**DOI:** 10.1101/2025.09.15.676443

**Authors:** Kelly X. Ji, Hongkai Ji

## Abstract

Spatially resolved assay for transposase-accessible chromatin with sequencing (spATAC-seq) is an emerging technology for studying spatial variation in gene regulatory landscapes within tissues. Current analysis pipelines commonly apply library size normalization, assuming that variation in sequencing library size across cells represents a technical confounder rather than biological signal. While recent studies have shown that library size can confound biological interpretation in spatial transcriptomics, its impact in spATAC-seq remains poorly understood. Here, we show that library size in spATAC-seq data is biologically informative and that standard normalization methods can obscure important biological signals and hinder downstream analyses. These findings underscore the need for caution and for the development of improved approaches to address library size in spATAC-seq analysis.

## Introduction

Advances in spatial omics technologies have enabled the systematic profiling of spatial patterns in cellular molecular landscapes. Among these, spatial RNA sequencing (spRNA-seq) has become widely used to investigate transcriptomes within their native tissue context.^1,2^ More recently, the development of spatial assay for transposase-accessible chromatin using sequencing (spATAC-seq) has made it possible to study the spatial organization of chromatin accessibility, thereby providing insights into the regulatory landscape of cells in situ.^3-6^ These technologies offer powerful tools for dissecting tissue architecture, exploring cellular responses to microenvironmental changes, and uncovering mechanisms of cell-cell interaction and communication, ultimately accelerating our understanding of both basic biology and disease.

In the analysis of spRNA-seq and spATAC-seq data, library size – typically defined as the total number of sequence reads or unique molecular identifiers (UMIs) per cell or spot – is commonly treated as a technical confounder.^4-7^ Standard workflows routinely normalize for library size to mitigate presumed technical variation before downstream analysis.^4-12^ However, a recent study has demonstrated that in spRNA-seq data, library size can reflect underlying tissue structure and that normalization may obscure biologically meaningful signals, adversely affecting spatial domain detection and interpretation.^13^ Whether a similar phenomenon exists in spATAC-seq data remains unclear. If library size in spATAC-seq also encodes biological information, current normalization practices could inadvertently remove relevant signals and hinder discovery.

To address this question, we analyzed five spATAC-seq datasets generated by three different laboratories using distinct technologies: spatial-ATAC-RNA-seq,^4^ spatial-Mux-seq,^5^ and Slide-tags^6^ (**Supplementary Table 1**). We observed multiple lines of evidence supporting that library size in spATAC-seq is indeed associated with biology.

## Results & Discussion

First, comparison of spatial maps of spATAC-seq library size with matched histology images revealed strong correlation between library size and tissue structure (**Fig. 1**). For instance, in the Human Hippocampus spatial-ATAC-RNA-seq dataset, the granule cell layer (GCL) exhibited markedly higher library size compared to surrounding regions (**Fig. 1a**). Clustering histology image pixels into five groups based on intensity (gray value) revealed that different histological clusters had significantly different mean library sizes (**Fig. 1b**), quantitatively confirming the association between library size and tissue architecture. Similar patterns were observed in the other four datasets (**Fig. 1c–j**). In the Human Melanoma dataset, spots were grouped into two clusters corresponding to the two tumor compartments annotated in the original study, whereas the remaining datasets used five clusters.

**Fig 1:**
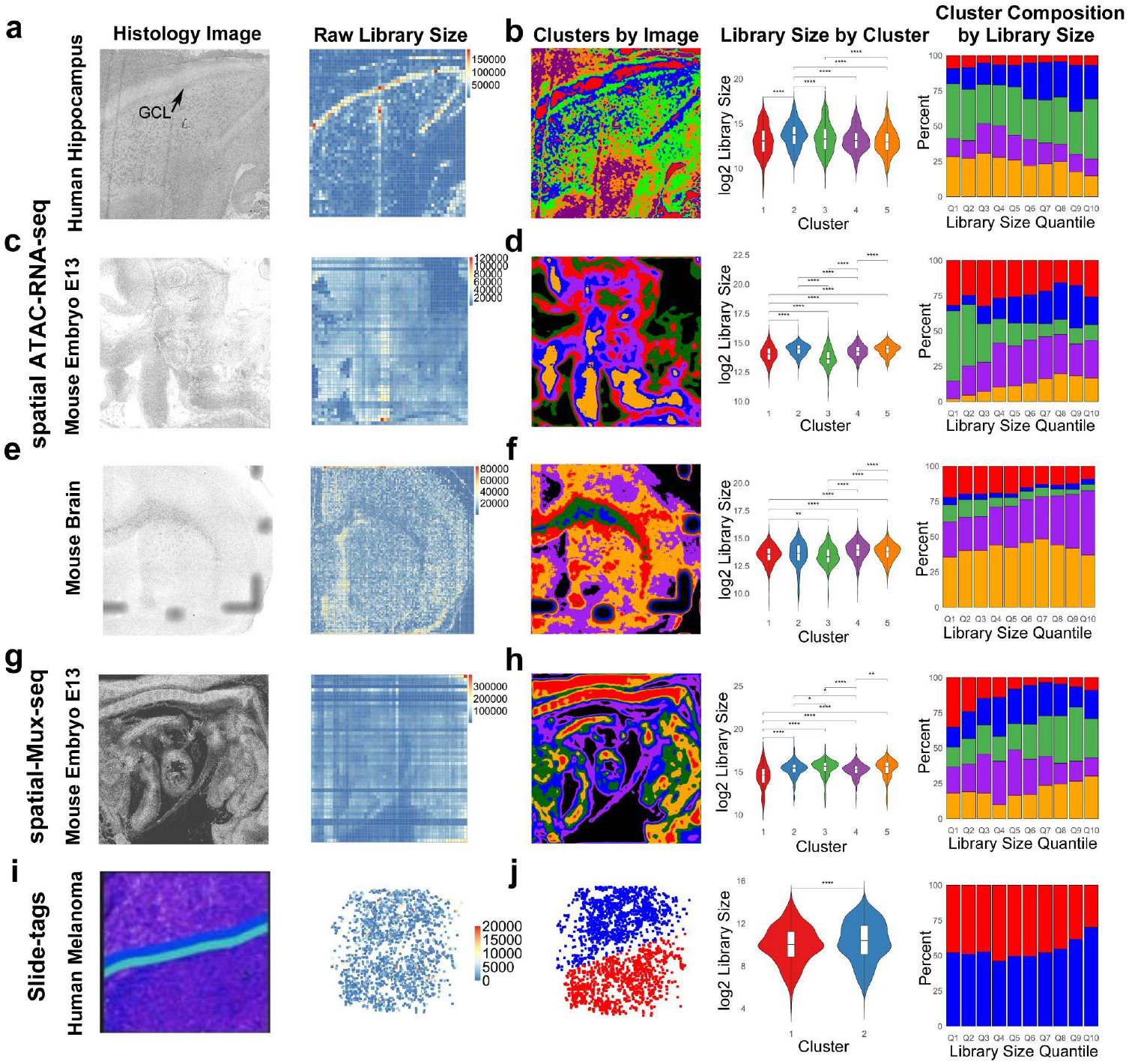
Library size is associated with tissue structure across spatial-ATAC datasets. a) Left - Raw histology images; Right - library size per spot of human hippocampus.^4^ b) Distribution and composition of library size across different histology-determined clusters in human hippocampus. From left to right: clusters of spots based on pixel intensity in histology image, distribution of spot-level library size across clusters, and composition of clusters by each 10% quantile of library size. Asterisks represent significance across two clusters calculated via a two-sided *t*-test (p-value<0.05 is *, p-value<0.01 is **, p-value<0.001 is ***, and p-value<0.0001 is ****). c) & d) Analysis on mouse embryo (E13).^4^ e) & f) Analysis on mouse brain.^4^ g) & h) Analysis on mouse embryo E13 spatial-Mux-seq dataset.^5^ i) & j) Analysis on human melanoma Slide-tags sequencing.^6^ The histology images^4,5^ (a, c, e, g) and the original annotated image^6^ (i) associated with the original publications^4-6^ are shown in the first column. Panel g: Histology image reproduced from Guo et al., *Nat Methods* 22:520–529 (2025)^5^, with permission from Springer Nature; Not covered by this manuscript’s CC-BY-NC license.

Second, we observed strong correlations between library size in spATAC-seq and library size in spatial RNA-seq for matched spatial spots across all datasets (**Fig. 2a**). To further investigate this relationship, we computed the correlation between promoter accessibility (ATAC-seq signal) and gene expression (RNA-seq signal) across spatial spots for each individual gene. Using raw RNA-seq count data as a benchmark, we found that gene expression correlated more strongly with raw (unnormalized) ATAC-seq signals than with library size-normalized ATAC-seq signals (**Fig. 2b**). Similar trends were observed when using library size-normalized RNA-seq data as the benchmark (**Fig. 2c**). Note that in these gene-level analyses, the overall correlation between RNA-seq and ATAC-seq was relatively low due to high levels of sparsity and noise of the data.

**Fig 2:**
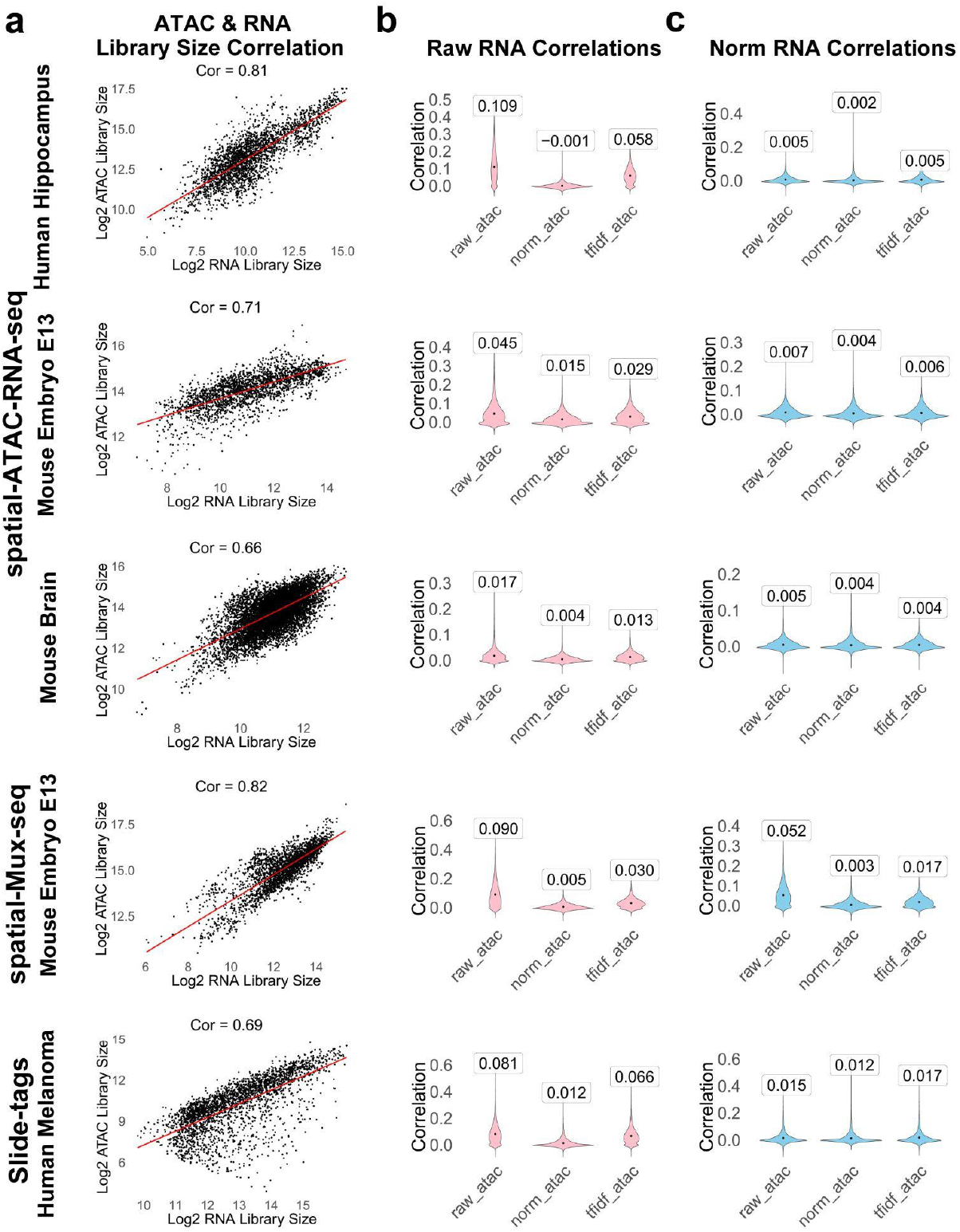
Correlation between ATAC-seq and RNA-seq. a) ATAC library size correlation with RNA library size across datasets. Each data point in the scatterplots is a spot from the spatial data. Cor: Pearson correlation. b) Raw, normalized, and TF-IDF corrected ATAC promoter activity correlations with raw RNA gene expression. c) Raw, normalized, and TF-IDF corrected ATAC promoter activity correlations with library-size-normalized RNA gene expression. In (b) and (c), each data point in the violin plot corresponds to a gene, ATAC reads in the ±5kb promoter region were summarized into the gene’s promoter activity, and the y-axis shows the Pearson correlation between the RNA gene expression and ATAC promoter activity across spatial spots. The values in the boxes on top of violin plots represent the mean Pearson correlation across all genes.

TF-IDF transformation is currently a state-of-the-art normalization method for single-cell and spatial ATAC-seq data,^8-12^ especially under conditions of extreme sparsity. However, even compared to TF-IDF–normalized ATAC-seq data, raw ATAC-seq signals exhibited stronger correlations with gene expression, regardless of whether raw or normalized RNA-seq was used (**Fig. 2b,c**). Using different window sizes (±5kb or ±2kb around transcription start sites) to summarize promoter ATAC signals yielded similar results (**Fig. 2, Supplementary Fig. 1**).

Notably, the strongest cross-modality correlation was observed when both RNA-seq and ATAC-seq data were left unnormalized (**Fig. 2b,c**). The gene-level concordance between raw RNA and raw ATAC signals may reflect either true biology (if library size carries biological meaning) or shared technical artifacts (if library size is purely a confounder). However, the observation that normalized RNA-seq (which removes the library size effect) still correlates more strongly with raw ATAC-seq than with normalized ATAC-seq suggests that biologically meaningful signals are being lost during ATAC-seq normalization. This supports the interpretation that library size in spATAC-seq is not merely a technical confounder but likely contains biological information.

Third, when comparing spatial domains detected from spATAC-seq data to those derived from histology images or manually annotated, the raw spATAC-seq data yielded the best performance, as indicated by the highest Adjusted Rand Index (ARI) in 4 of 5 datasets (**Fig. 3**). It outperformed both library-size–normalized spATAC-seq data and TF-IDF-transformed data. In the human melanoma dataset, the improvement was substantial: domains identified from raw spATAC-seq data clearly separated two tumor compartments with distinct DNA copy number variation (CNV) profiles (ARI = 0.90), whereas neither library size normalization (ARI=0.14) nor TF-IDF (ARI=0.0009) was able to distinguish these populations (**Fig. 3e**). Performing this analysis with different numbers of spatial domains (*K* = 5 or *K* = 10) shows that the overall performance of raw data is consistently better than or comparable to that of the normalized data, indicating the robustness of this conclusion (**Fig. 3, Supplementary Fig. 2**).

**Fig 3:**
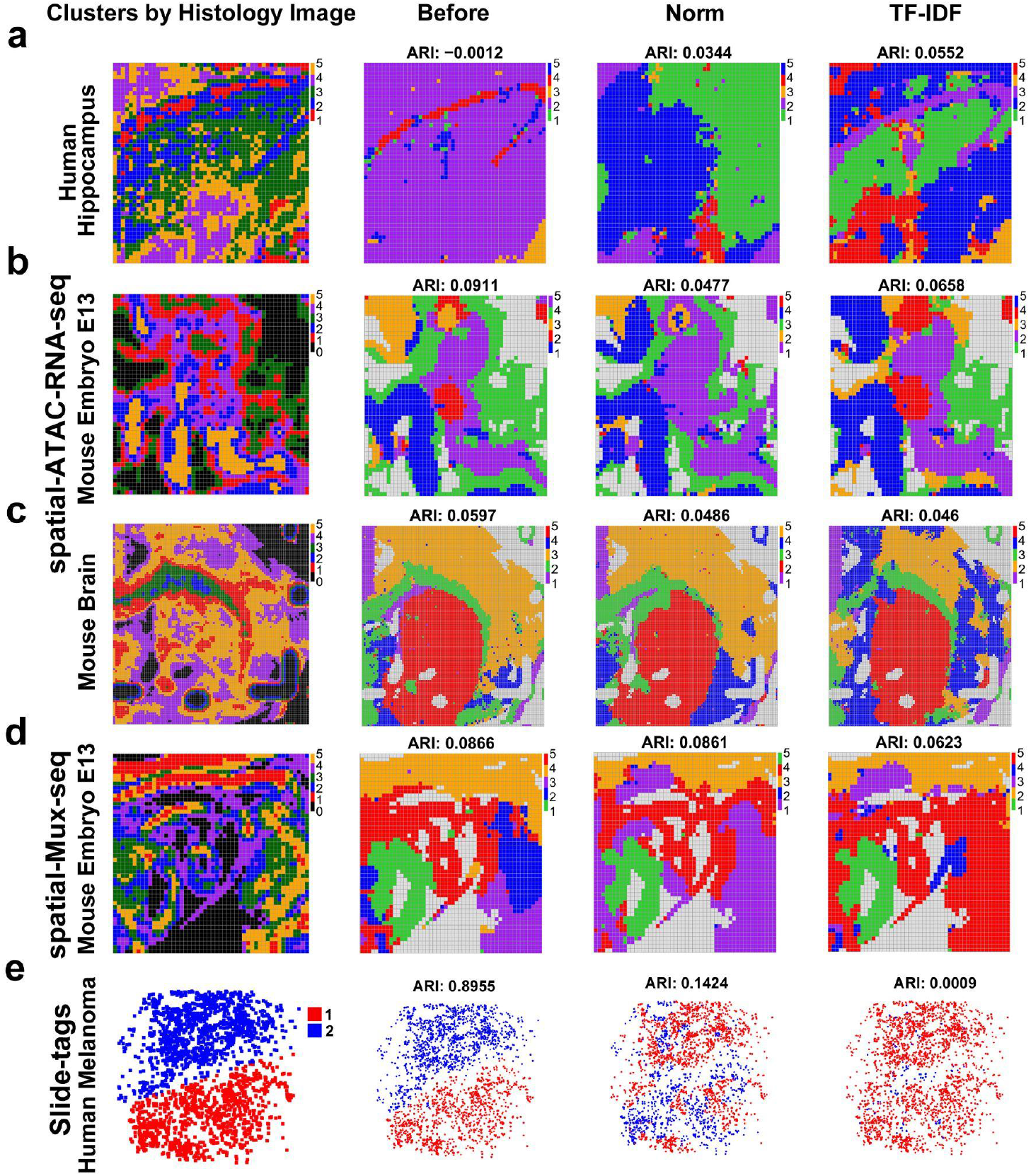
Spatial domain detection performance before versus after library size normalization. a) Human hippocampus spatial-ATAC-RNA data. b) Mouse embryo spatial-ATAC-RNA data. c) Mouse brain spatial-ATAC-RNA data. d) Mouse embryo spatial-Mux-seq data. e) Human melanoma Slide-tags data. From left to right: spatial clusters based on k-means clustering on histology image pixel intensities (a-d) or spatial domain annotations from the original study based on copy number variation profiles (e), spatial domains detected by BASS using raw ATAC counts, library-size-normalized counts, and TF-IDF values. The number of spatial clusters/domains is set to *K*=5 for panels (a-d) and *K*=2 for panel (e). ARIs are shown on top.

Collectively, the above results suggest that library size is associated with underlying biological variation. As such, standard library size normalization, though widely used, may inadvertently remove meaningful biological signals. This can weaken the correlation between ATAC-seq and RNA-seq data and impair the detection of spatial domains.

Importantly, whether or not normalization is applied can lead to different conclusions when identifying genomic loci with differential signals across spatial spots or domains. For example, when comparing the GCL domain with other spatial domains in the human hippocampus dataset, 10,876 differential promoters were identified by both raw and library size–normalized spATAC-seq data using Wilcoxon rank-sum tests (FDR≤0.05). Among these, 9805 (90.2%) exhibited opposite directions of change (e.g., a promoter upregulated in GCL according to the normalized data appeared downregulated in the raw data) (**Fig. 4a,b**).

**Fig 4:**
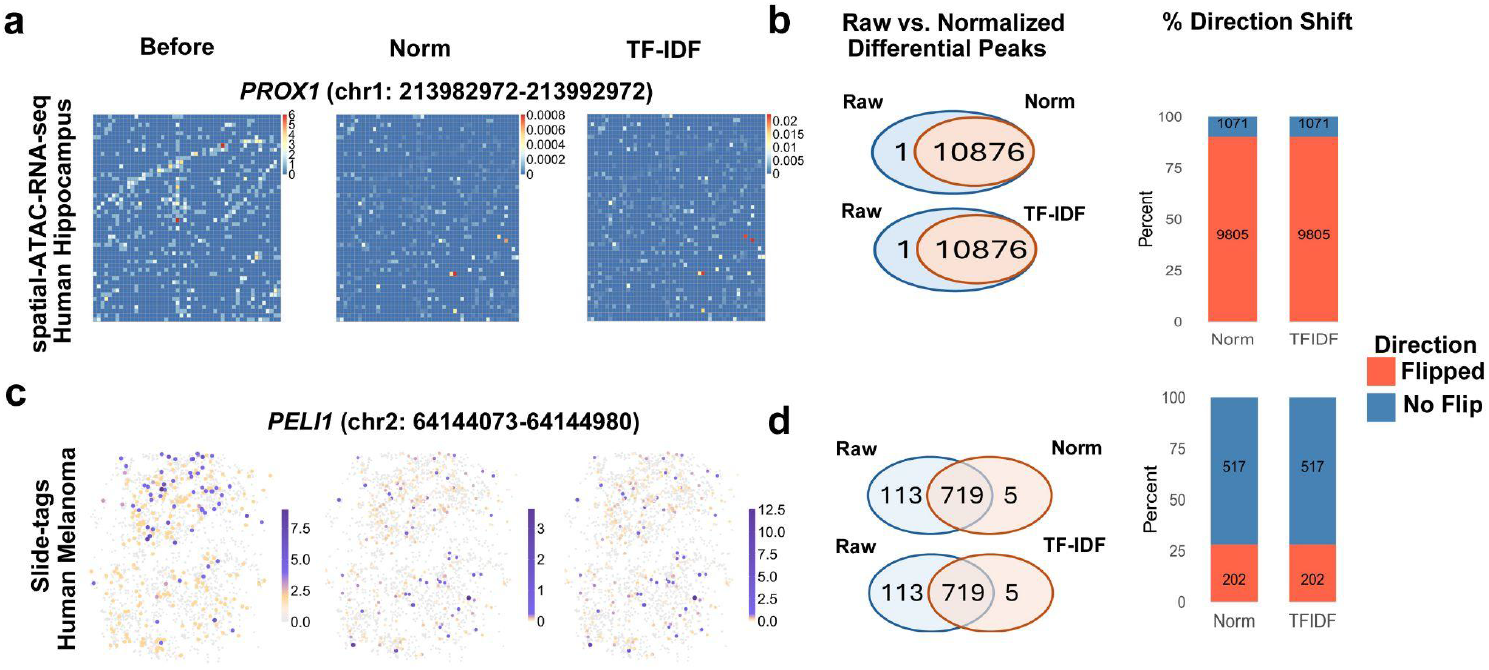
Differential promoter activity shifts direction after library size normalization. a) Example on human hippocampus dataset. From left to right: ATAC promoter activity for *PROX1* (chr1:213982972-213992972) characterized using raw count, library-size normalized count, and TF-IDF transformed data. *PROX1* is known to be upregulated in the GCL region.^14^ b) The venn diagram shows the overlap between raw vs. normalized and TF-IDF differential promoters in the human hippocampus dataset. For the 10,876 differential promoters detected by both raw and normalized counts, the stacked barplot shows the percentage of promoters that switched the directions (orange) and the percentage with concordant directions (blue). c) Example on Slide-tags human melanoma dataset. ATAC peak activity for the *PELI1* gene is shown. d) Similar to b) but for the human melanoma dataset.

Comparing differential promoter analysis results using raw counts and TF-IDF-transformed data yielded identical findings (**Fig. 4a,b**). TF-IDF combines term frequency with inverse document frequency, where the term frequency corresponds to the library-size-normalized count. For a given promoter, while the term frequency may vary across spots, the inverse document frequency is constant across all spots. As a result, the TF-IDF values for any given promoter represent a monotonic transformation of its normalized counts. Since the Wilcoxon rank-sum test used for differential promoter detection is rank-based and thus invariant to monotonic transformations, the sets of differential promoters identified using normalized counts and TF-IDF values are expected to be identical.

As a concrete example, **Fig. 4a** shows the spatial activity of a promoter corresponding to the *PROX1 gene* (chr1:213982972-213992972) before and after normalization. *PROX1* is known to have upregulated gene expression in the GCL region.^14^ However, the upregulated promoter activity in the GCL region before normalization disappeared after library size normalization or TF-IDF transformation.

Similarly, in the human melanoma dataset, comparing two tumor compartments revealed 719 differential regions detected by both approaches, among which 202 (28%) showed opposite directions. These discrepancies underscore that library size normalization can substantially alter biological interpretations, even reversing conclusions for many genomic loci (**Fig. 4c,d**).

These findings highlight the need for caution in handling library size variation in spATAC-seq data. Because library size may reflect both meaningful biological differences and unwanted technical variation, and the two are often perfectly confounded, this presents a major analytical challenge. Resolving this confounding remains an open problem and a critical unmet need.

Addressing it, whether through improved experimental or computational methods, is essential for fully realizing the potential of spATAC-seq in spatial epigenomics. Finally, the compiled data and evaluation pipeline from our study provide a valuable resource for benchmarking future computational methods aimed at tackling this issue.

## Methods

### Data Collection

Five publicly available spatial ATAC-seq datasets, along with co-profiled spatial RNA-seq, were analyzed in this study. Spatial-ATAC-RNA-seq coprofiling datasets from Human Hippocampus (at 50 µm resolution), Mouse Embryo (50 µm), and Mouse Brain (20 µm) were downloaded from the Gene Expression Omnibus (GEO Accession: GSE205055). The Mouse Embryo spatial-Mux-seq dataset (50 µm) generated using multiplexed spatial mapping was obtained from GEO (Accession: GSE263333). The Human Melanoma Slide-tags dataset was downloaded from the Broad Institute Single Cell Portal (Accession: SCP2176). All datasets included either original histology or annotated images, spatial coordinates, and ATAC- and RNA-seq count matrices generated by aligning FASTQ files to the reference genomes (mm10 for mouse and hg38 for human). We downloaded aligned TSV matrices and corresponding spatial metadata. All downstream analyses were conducted in R (version 4.5.1).

### Histology Image Processing and Clustering

For spatial-ATAC-RNA-seq and spatial-Mux-seq datasets, raw H&E-stained brightfield histology images were first smoothed using average pixel intensity and a sliding window (51-pixel x 51-pixel). K-means clustering was then applied to segment the images by grouping pixels into *K* (=5 by default) clusters based on pixel intensity. For the Slide-tags human melanoma dataset, the original publication identified two tumor compartments based on distinct copy number variation profiles. These two compartments (*K*=2) were used as two ground truth clusters.

After clustering pixels, the resolution of each spatial-ATAC-RNA-seq and spatial-Mux-seq histology image was downsampled to match the spatial resolution of the spot data in spATAC-seq. Downsampling was performed by dividing the image into blocks. This yielded 50×50 or 100×100 matrices corresponding to the spatial layout of each spATAC-seq dataset. Each spot was assigned a cluster label based on the most abundant cluster label of the pixels within the block. Differences in library size between clusters were evaluated using two-sample t-tests.

### Spatial ATAC-seq Data Processing

To mitigate data sparsity, spatial ATAC-seq data were summarized based on promoter activity. RefSeq gene annotations for mm10 and hg38 genomes were downloaded from the UCSC Genome Browser.^15^ Promoter regions were defined as ±2 kb and ±5 kb around genes’ transcription start sites (TSS). Duplicated entries were removed, retaining one transcript per gene symbol. To create the promoter activities, ATAC fragments overlapping promoter regions were counted using the Genomic Ranges R package (v1.60.0),^16^ producing promoter-by-spot matrices.

### Normalization Methods for Benchmarking

To benchmark the impact of library size normalization, three methods were compared, including raw count without normalization, library size normalization, and Term Frequency-Inverse Document Frequency (TF-IDF).^17^ For library size normalization, each raw count was divided by the corresponding library size and multiplied by a scale factor of 10,000. For TF-IDF, which is the state-of-the-art technique for single cell ATAC-sequencing datasets, default parameters for the RunTFIDF function in Signac was used to convert raw counts into TF-IDF values.^9^

### Correlation with RNA

First, we evaluated the Pearson correlation between the library sizes of spatial RNA and spatial ATAC datasets using the co-profiling data.

Next, we compared no normalization, library size normalization and TF-IDF by evaluating the Pearson correlation of each gene’s transcription level (RNA-seq) and promoter activity (ATAC-seq) across spatial spots. Gene’s transcription level was summarized using either raw RNA-seq counts without normalization or library size normalized counts. Promoter activity was summarized using either raw counts, library-size normalized counts, or TF-IDF values. The distribution of correlation across all genes, along with the mean correlation are shown as violin plots in Fig. 2.

### Spatial Domain Detection

Spatial domain detection was performed using either ATAC promoter activity (defined as ±5 kb from a gene’s transcription start site) for spatial-ATAC-RNA-seq and spatial-Mux-seq or peak regions provided in the original paper for Slide-tags data. BASS (Bayesian Analytics for Spatial Segmentation)^18^ was used to group spatial spots into *K* distinct domains (*K*=2 for Slide-tags and *K*=5 and 10 for other datasets; cell type number was set to the default value 15 for all datasets). Correspondingly, pixels in the histology image were also segmented into *K* clusters based on the pixel intensity. Adjusted rand index (ARI) was calculated between spATAC-seq identified domains and histology image identified clusters. The ARIs from different normalization methods are then compared. Higher ARI values indicate better performance.

BASS is originally developed for spatial domain detection using spatial transcriptomic data,^18^ with improved performance when benchmarked against other high-performing spatial domain detection tools.^19^ Here it is applied to 30 top principal components (PCs) of the promoter or peak activity data to detect domains in spATAC-seq data. For other parameters, BASS was run using their default values.

### Differential Promoter Activity Analysis

To further explore the impact of library size normalization on downstream analysis, we conducted differential promoter analysis using two datasets with prior annotations: the human melanoma data with previously annotated two tumor compartments, and the human hippocampus data with a clear GCL domain. For the human hippocampus spatial-ATAC-RNA-seq data, promoters were defined as ±5kb from a gene’s transcription start site, and we compared spots within and outside the GCL domain (as defined in the original paper as cluster 4) to detect differential signals. For the human melanoma Slide-tags data, peaks downloaded from the original study were used. Promoters or peaks with differential activities were identified using Wilcoxon rank-sum tests. Two-sided p-values were converted into false discovery rate (FDR) using the Benjamini-Hochberg procedure to correct for multiple testing.^20^ The analysis was run using raw count data, library-size normalized data, or TF-IDF transformed data, respectively.

FDR≤0.05 was used to call differential promoter or peak regions. We then counted the differential regions in each analysis and compared the results before normalization (raw counts) and after normalization (library-size normalization, TF-IDF). The numbers of common and method-specific differential regions were summarized into venn diagrams. Next, among the common differential regions detected by both methods, we also calculated the percentage of differential regions that switched the direction (e.g., reported as upregulated before normalization but downregulated after normalization, or reported as downregulated before normalization but upregulated after normalization) and displayed the results in stacked barplots.

## Resource Availability

### Code Availability

All code used to produce the analyses reported in this paper can be accessed on GitHub via the following link: github.com/tulipblossoms/spATAC-Library-Size

### Data Availability

Five datasets were used for this study. The 50 μm Human Hippocampus, 50 μm Mouse Embryo, and 20 μm Mouse Brain spatial-ATAC-RNA seq coprofiling datasets can be downloaded from the Gene Expression Omnibus (GEO Accession: GSE205055). 50 μm Mouse Embryo spatial-Mux-seq data can be downloaded from Gene Expression Omnibus (GEO Accession: GSE263333). Human Melanoma Slide-tags data can be downloaded from the Broad Institute Single Cell Portal (Accession Number: SCP2176). Annotations for genes (file names mm10.refGene and hg38.refGene) can be acquired from the UCSC Genome Browser.

## Acknowledgements

We would like to thank the editor and reviewers for their valuable feedback.

## Author Contributions

Conceptualization, H.J.; Methodology, H.J. & K.J.; Formal Analysis, K.J.; Visualization, K.J.; Writing-Original Draft, K.J. and H.J.; Writing-Review & Editing, K.J. and H.J.

## References

1. Marx, V. Method of the Year: spatially resolved transcriptomics. Nat Methods 18, 9–14 (2021). 10.1038/s41592-020-01033-y

2. Crosetto, N., Bienko, M. & van Oudenaarden, A. Spatially resolved transcriptomics and beyond. Nat Rev Genet 16, 57–66 (2015). 10.1038/nrg3832

3. Deng, Y., Bartosovic, M., Ma, S. et al. Spatial profiling of chromatin accessibility in mouse and human tissues. Nature 609, 375–383 (2022). 10.1038/s41586-022-05094-1

4. Zhang, D., Deng, Y., Kukanja, P. et al. Spatial epigenome–transcriptome co-profiling of mammalian tissues. Nature 616, 113–122 (2023). 10.1038/s41586-023-05795-1

5. Guo, P., Mao, L., Chen, Y. et al. Multiplexed spatial mapping of chromatin features, transcriptome and proteins in tissues. Nat Methods 22, 520–529 (2025). 10.1038/s41592-024-02576-0

6. Russell, A.J.C., Weir, J.A., Nadaf, N.M. et al. Slide-tags enables single-nucleus barcoding for multimodal spatial genomics. Nature 625, 101–109 (2024). 10.1038/s41586-023-06837-4

7. Luecken MD, Theis FJ. Current best practices in single-cell RNA-seq analysis: a tutorial. Mol Syst Biol. 2019;15:e8746.

8. Chen, X., Li, K., Wu, X. et al. Descart: a method for detecting spatial chromatin accessibility patterns with inter-cellular correlations. Genome Biol 25, 322 (2024). 10.1186/s13059-024-03458-6

9. Stuart, T., Srivastava, A., Madad, S. et al. Single-cell chromatin state analysis with Signac. Nat Methods 18, 1333–1341 (2021). 10.1038/s41592-021-01282-5

10. Butler, A., Hoffman, P., Smibert, P. et al. Integrating single-cell transcriptomic data across different conditions, technologies, and species. Nat Biotechnol 36, 411–420 (2018). 10.1038/nbt.4096

11. Cao, X., Ma, T., Fan, R., Yuan, G. C. (2024). Systematic analysis identifies a connection between spatial and genomic variations of chromatin states. Cell Systems, 15(11), 1092–1102.e2. 10.1016/j.cels.2024.10.006

12. Granja, J.M., Corces, M.R., Pierce, S.E. et al. ArchR is a scalable software package for integrative single-cell chromatin accessibility analysis. Nat Genet 53, 403–411 (2021). 10.1038/s41588-021-00790-6

13. Bhuva, D.D., Tan, C.W., Salim, A. et al. Library size confounds biology in spatial transcriptomics data. Genome Biol 25, 99 (2024). 10.1186/s13059-024-03241-7

14. Lavado, A., Lagutin, O. V., Chow, L. M., Baker, S. J., Oliver, G. (2010). Prox1 is required for granule cell maturation and intermediate progenitor maintenance during brain neurogenesis. PLoS Biol, 8(8), e1000460. 10.1371/journal.pbio.1000460

15. Perez, G., Barber, G. P., Benet-Pages, A., Casper, J., Clawson, H., Diekhans, M., Fischer, C., Gonzalez, J. N., Hinrichs, A. S., Lee, C. M., Nassar, L. R., Raney, B. J., Speir, M. L., van Baren, M. J., Vaske, C. J., Haussler, D., Kent, W. J., Haeussler, M. (2025). The UCSC Genome Browser database: 2025 update. Nucleic Acids Res, 53(D1), D1243–D1249. 10.1093/nar/gkae974

16. Lawrence M, Huber W, Pagès H, Aboyoun P, Carlson M, Gentleman R, et al. (2013) Software for Computing and Annotating Genomic Ranges. PLoS Comput Biol 9(8): e1003118. 10.1371/journal.pcbi.1003118

17. Spärck Jones, K. (1972). A statistical interpretation of term specificity and its application in retrieval. Journal of Documentation, 28(1), 11–21. 10.1108/eb026526

18. Li, Z., Zhou, X. BASS: multi-scale and multi-sample analysis enables accurate cell type clustering and spatial domain detection in spatial transcriptomic studies. Genome Biol 23, 168 (2022). 10.1186/s13059-022-02734-7

19. Yuan, Z., Zhao, F., Lin, S. et al. Benchmarking spatial clustering methods with spatially resolved transcriptomics data. Nat Methods 21, 712–722 (2024). 10.1038/s41592-024-02215-8

20. Benjamini, Y., Hochberg, Y. (1995). Controlling the False Discovery Rate: A Practical and Powerful Approach to Multiple Testing. Journal of the Royal Statistical Society. Series B (Methodological), 57(1), 289–300. http://www.jstor.org/stable/2346101

