## Supplementary Table 1, Supplementary Fig. 1, Supplementary Fig. 2 for "Library Size in Spatial ATAC-seq: Technical Confounder or Biology?"

### Supplementary Materials

**Supplementary Table 1**: 5 spatial multi-modal (i.e., ATAC + RNA) datasets across 2 species, 2 resolutions, and 3 different technologies were used to study the impact of library size normalization on biology in spatial-ATAC sequencing. The dataset accession number, species, tissue-type, resolution, and technology of the 5 datasets included in this study are listed below.

| **Dataset Accession** | **Species** | **Tissue Type** | **Resolution** | **Number of Spots** | **Technology** |
|---|---|---|---|---|---|
| GSE205055 | Human | Hippocampus | 50 µm | 2500 | spatial-ATAC-RNA-seq |
| GSE205055 | Mouse | Embryo E13 | 50 µm | 2500 | spatial-ATAC-RNA-seq |
| GSE205055 | Mouse | Brain | 20 µm | 10000 | spatial-ATAC-RNA-seq |
| GSE263333 | Mouse | Embryo E13 | 50 µm | 2500 | spatial-Mux-seq |
| SCP2176 in Broad Single Cell Portal | Human | Melanoma | 20 µm | 2535 | Slide-tags |

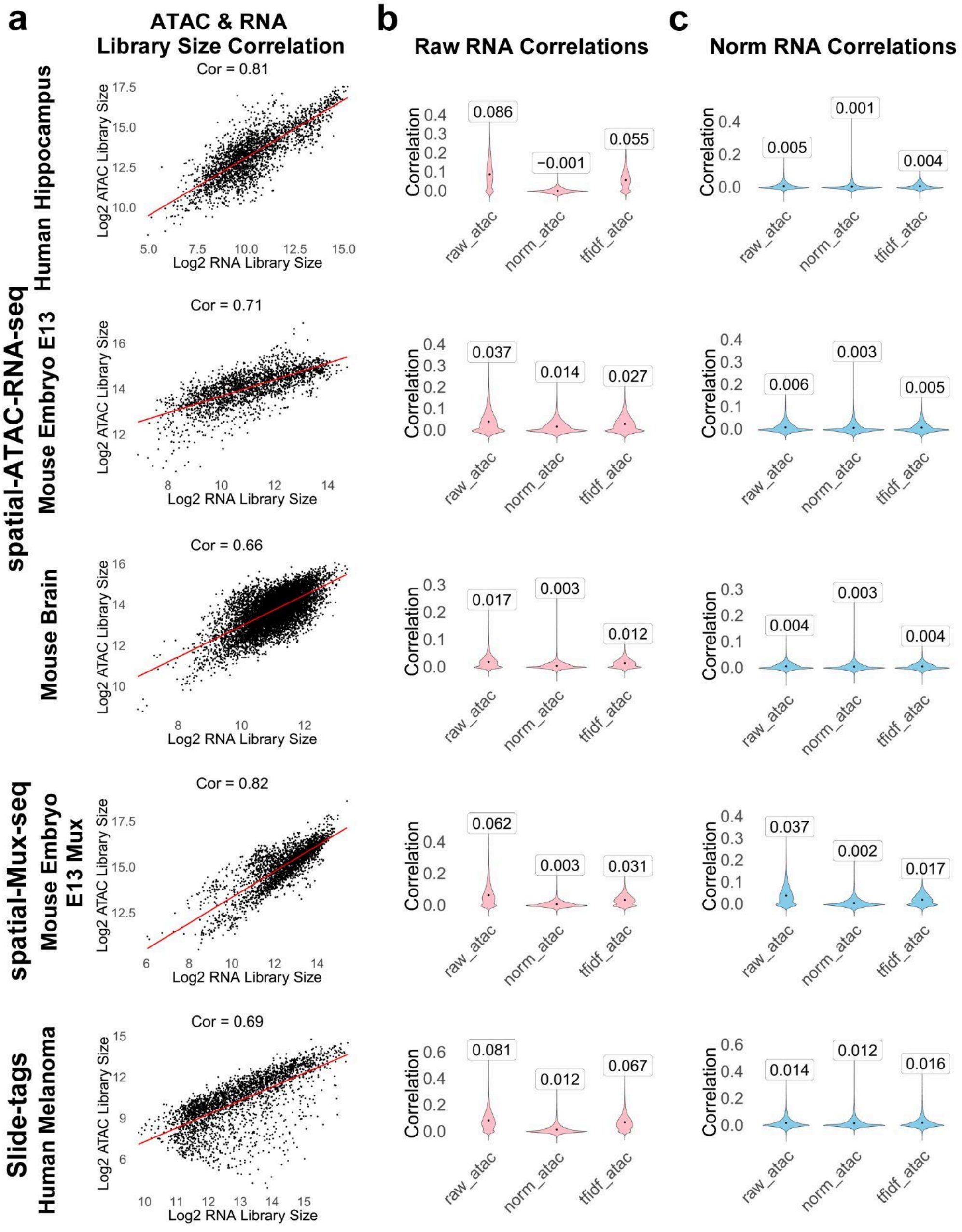


**Supplementary Fig 1**: Correlation between ATAC-seq and RNA-seq. a) ATAC library size correlation with RNA library size across datasets. Each data point in the scatterplots is a spot from the spatial data. Cor: Pearson correlation. b) Raw, normalized, and TF-IDF corrected ATAC promoter activity correlations with raw RNA gene expression. c) Raw, normalized, and TF-IDF corrected ATAC promoter activity correlations with library-size-normalized RNA gene expression. In (b) and (c), each data point in the violin plot corresponds to a gene, ATAC reads in the ±2kb promoter region were summarized into the gene's promoter activity, and the y-axis shows the Pearson correlation between the RNA gene expression and ATAC promoter activity across spatial spots. The values in the boxes on top of the violin plots represent the mean Pearson correlation across all genes.

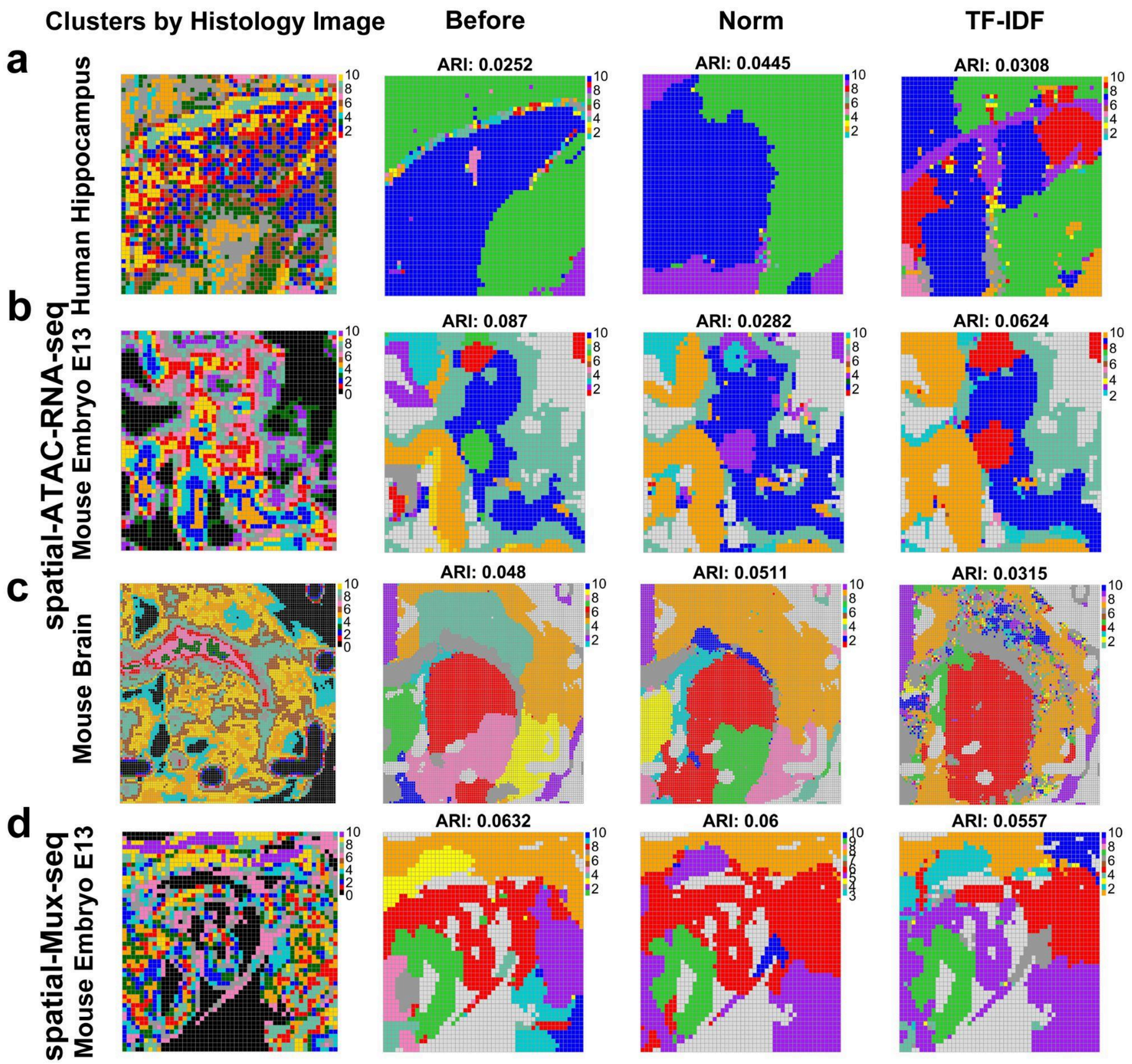


**Supplementary Fig 2**: Spatial domain detection performance before versus after library size normalization with $K$=10 domains. a) Human hippocampus spatial-ATAC-RNA data. b) Mouse embryo spatial-ATAC-RNA data. c) Mouse brain spatial-ATAC-RNA data. d) Mouse embryo spatial-Mux-seq data. From left to right: spatial clusters based on k-means clustering on histology image pixel intensities (a-d), spatial domains detected by BASS using raw ATAC counts, library-size-normalized counts, and TF-IDF values. ARIs are shown on top.
